# Modulatory effects of α7-nicotinic cholinergic receptors on perceptual sensitivity in a visual signal detection task

**DOI:** 10.64898/2026.05.18.725386

**Authors:** Harry J. Robson, Alexander R.H. Matthews, Livia J.F. Wilod Versprille, Johann F. du Hoffmann, Jeffrey W. Dalley

**Affiliations:** Department of Psychology, University of Cambridge, Downing St, Cambridge CB2 3EB, UK; Department of Systems Neuroscience, Johann-Friedrich-Blumenbach Institut für Zoologie und Anthropologie, Universität Göttingen, Göttingen, Germany; Boehringer Ingelheim Pharma GmbH & Co. KG, Div. Research Germany, Biberach an der Riß, Germany; Department of Psychiatry, Hershel Smith Building for Brain and Mind Sciences, University of Cambridge, Cambridge CB2 0SZ, UK

**Keywords:** acetylcholine, visual attention, nicotinic receptors, signal detection theory

## Abstract

**Rationale:** Cholinergic signalling is critical for attentional control and signal detection, yet the contribution of specific acetylcholine receptor (AChR) subtypes remains poorly understood. Although the α7 nicotinic AChR (nAChR) holds promise as a target for cognition-enhancing therapy, clinical findings to date have been inconsistent.

**Objective:** To investigate the effects of putative cognitive enhancing drugs, including those targeting cholinergic transmission and α7 nAChRs on a visual signal detection task (SDT).

**Methods:** Male and female Sprague Dawley rats were trained on an SDT. Cholinergic transmission was probed systemically with nicotinic and muscarinic receptor antagonists (mecamylamine and scopolamine), a cholinesterase inhibitor (galantamine), an M4-AChR positive allosteric modulator (PAM; VU0467154), an α7 nAChR antagonist (MLA), an α7 nAChR PAM (CCMI), and an α7 nAChR partial agonist (SSR-180,711). Dopaminergic transmission was probed using the catechol-O-methyltransferase (COMT) inhibitor, tolcapone. A novel, trial-level signal detection theory-based generalised linear mixed-effects model (SDT-GLMM) was used to index response bias and perceptual sensitivity (d′), the latter reflecting subjects’ ability to discriminate signal from noise.

**Results:** Mecamylamine profoundly impaired SDT performance across all measures. Galantamine significantly improved d′ at moderate doses but not when a distractor was present. MLA uniquely produced dose-dependent improvements in d′ that were preserved under distraction. In contrast, positive allosteric modulation and agonism of α7 nAChRs impaired task performance. Scopolamine, VU0467154, and tolcapone had no consistent or interpretable effects on signal detection.

**Conclusions:** This work demonstrates that α7 nAChR modulation bidirectionally and dose-dependently regulates perceptual sensitivity, irrespective of attentional distraction. These findings have implications for targeted cognitive enhancement in disorders of attention.

## Introduction

Failures of attention are among the earliest and most debilitating features of many brain disorders, including Alzheimer’s disease (Chiu et al. 2004), attention-deficit / hyperactivity disorder (ADHD) (Fleming et al. 2017), and the cognitive dysfunction present in schizophrenia and Parkinson’s disease (Braff 1993; Brown and Marsden 1988). Converging evidence implicates cholinergic dysfunction in these disorders (Whitehouse et al. 1981; Freedman et al. 1995; Sarter and Bruno 2004), yet current cholinergic treatments, while clinically effective for memory symptoms, provide only modest attentional benefits and often result in significant adverse side effects (Pepeu et al. 2013; Marucci et al. 2021). A more precise mechanistic understanding of how the cholinergic systems contribute to cognitive dysfunction is thus needed to develop more effective treatments.

To optimise action selection and guide behaviour in noisy, uncertain environments, the brain must prioritise limited processing resources to attentionally relevant inputs (Posner et al. 1980; Hasselmo and Sarter 2011; Dehaene and Changeux 2011). Signal detection tasks (SDTs) provide a tractable framework for studying such attentional processes, by requiring subjects to distinguish signal-presence from signal-absence while maintaining appropriate response control (Turner et al. 2016; Wilod Versprille et al. 2026). Within this framework, we operationalise a key component of attention, perceptual sensitivity (d′), which isolates cue discrimination from response bias (Tanner and Swets 1954; Bashinski and Bacharach 1980). Despite this advantage, many preclinical studies continue to analyse task performance using summary measures such as percentage correct, limiting mechanistic interpretation. Furthermore, while some studies appropriately quantify performance using d′ and criterion, these are often derived from aggregated data and analysed using ANOVA or linear mixed-effects models. This aggregation collapses trial-by-trial variability whereas modelling trial-level outcomes within a GLMM preserves the binomial structure of the data, avoids information loss due to averaging, and enables an improved dissociation of drug effects on d′ and response bias across experimental covariates (de Melo et al. 2022; DeCarlo 1998).

Cortical cholinergic signalling is a key determinant of signal detection, particularly under conditions of uncertainty (Sarter et al. 2005; Hasselmo and Sarter 2011; Sarter et al. 2014; Ballinger et al. 2016). In rodents performing signal detection, detected cues are associated with brief, phasic increases in prefrontal acetylcholine (ACh) release, whereas missed and non-cued trials lack such transients (Parikh et al. 2007; Sarter et al. 2014; Gritton et al. 2016). Phasic signals are thought to support the transition from monitoring to cue-directed responding by enhancing cortical signal-to-noise and enabling weak but behaviourally relevant stimuli to guide behaviour (Sarter et al. 2009; Hasselmo and Sarter 2011; Sarter and Lustig 2020). In contrast, tonic changes in extracellular ACh are associated with attentional effort (Kozak et al. 2006; Parikh et al. 2007), which notably occur with the presentation of distractors where perceptual uncertainty is increased (Himmelheber et al. 2000; Passetti et al. 2000; Sarter et al. 2006).

Nicotinic acetylcholine receptors (nAChRs) are important mediators of cholinergic effects on attention and detection (Sarter et al. 2014; Parikh and Bangasser 2020). The α7 nAChR has received particular attention as a candidate mediator of attentional control (Wallace and Bertrand 2013). α7 nAChRs are widely expressed in regions implicated in attention, including the prefrontal cortex, hippocampus, and basal forebrain. They are located pre- and post-synaptically on glutamatergic and GABAergic elements, with downstream influence on local circuit dynamics and dopaminergic signalling (Dani and Bertrand 2007; Livingstone et al. 2009; Koukouli and Maskos 2015). The high calcium permeability and rapid desensitisation kinetics of α7 nAChRs allow them to shape neurotransmission on behaviourally relevant timescales (Revah et al. 1991; Dani and Bertrand 2007; Albuquerque et al. 2009). Importantly, α7 nAChRs have been implicated in controlling the duration, but not the amplitude, of cue-evoked cholinergic transients in the medial prefrontal cortex (Parikh et al. 2010), positioning them as potential regulators of the temporally precise phasic cholinergic signalling required for reliable signal detection.

Pharmacological manipulation of α7 nAChRs has yielded complex and often dose-dependent effects on cognition. While α7 nAChR agonists and positive allosteric modulators can improve attentional performance in preclinical models (e.g., Wallace et al. 2011; Nikiforuk et al. 2016), clinical trials have produced mixed results (Freedman 2014), potentially reflecting rapid receptor desensitisation and narrow optimal dosing ranges (Revah et al. 1991; Cools and Arnsten 2022). Consistent with this interpretation, low doses of the selective α7 nAChR antagonist methyllycaconitine citrate (MLA) are reported to improve attentional performance in rodents (Hahn et al. 2011; Burke et al. 2014), suggesting that both insufficient and excessive α7 nAChR-related signalling may impair performance depending on baseline state and task demands. These findings raise the possibility that constraining α7 nAChR-mediated signalling may, under certain conditions, enhance signal detection processing.

The present study was designed to characterise how cholinergic interventions influence signal detection sensitivity, with a particular focus on α7 nAChRs under baseline and attentionally demanding conditions. Using a modified signal detection task (Turner et al. 2016), we investigated the effects of several compounds: a nicotinic receptor antagonist (mecamylamine), a muscarinic receptor antagonist (scopolamine), an M4-AChR positive allosteric modulator (PAM; VU0467154) an acetylcholinesterase inhibitor widely prescribed for the treatment of mild to moderate AD (galantamine; Marucci et al. 2021), a selective α7 nAChR antagonist (MLA), an α7 nAChR PAM (CCMI) and partial agonist (SSR-180,711), and a catechol-O-methyltransferase (COMT) inhibitor (tolcapone). Perceptual sensitivity (d′) was the primary outcome measure, reflecting the ability to discriminate signal presence from absence under uncertainty (Sarter et al. 2014). We hypothesised that: (1) mecamylamine would impair d′, since nicotinic cholinergic mechanisms support successful cue detection (Sarter et al. 2009; Hasselmo and Sarter 2011); (2) scopolamine and VU0467154 would have limited effects on d′, reflecting a lack of muscarinic receptor involvement; (3) MLA would produce dose-dependent biphasic effects consistent with α7 nAChR involvement in shaping cue-evoked signalling dynamics (Parikh et al. 2010; Hahn et al. 2011); (4) galantamine would exert dose-dependent effects confined to a narrow dose range and attenuated under distraction, consistent with a limitation of non-selective cholinergic enhancement for improving phasic detection mechanisms; (5) α7 activation would produce bidirectional, dose-dependent effects on SDT performance; (6) COMT inhibition would improve task performance, in line with previous work showing dopaminergic involvement in the SDT, and the positive relationship between α7 nAChRs and downstream dopaminergic signalling (Wilod Versprille et al. 2026; Livingstone et al. 2009). Critically, this approach enables identification of bidirectional and context-dependent effects of α7 nAChR modulation on d′ at the trial level.

## Methods

### Animals

Two cohorts of animals were used for this study. In each cohort, subjects were 32 Sprague-Dawley rats (16 females; Charles River, UK). All rats were approximately eight weeks old at the beginning of training, males weighed between 200-250g, and females weighed between 150-200g. In the first cohort, one female subject was removed from the statistical analyses as it did not receive all pharmacological agents. The animals were initially housed in cages of four in an animal facility at a constant temperature of 21°C and on a reversed 12h light-dark cycle (lights on at 0700). After reaching 400g, the males were separated into cages of two. Environmental enrichment was provided in the form of wooden chew blocks and cardboard tubes. The animals were food restricted to maintain weight at 85-95% of free-feeding weight, and given *ad libitum* access to water, except when undergoing behavioural procedures. All experimental sessions took place between 1000-1400 each day. All animal procedures were regulated in accordance with the Animals (Scientific Procedures) Act 1986 Amendment Regulations 2012, following the ethical approval by the University of Cambridge Animal Welfare and Ethical Review Body (AWERB). Experimental procedures were conducted under the project licenses of Prof. Amy Milton (PPLs PA9FBFA9F, PP9536688).

### Behavioural apparatus and protocol

A schematic of the operant chamber used for the SDT can be seen in Fig 1. For each trial, following an inter-trial interval (ITI), the central nose poke is illuminated, allowing the rat to make a nose poke to begin the trial. Immediately following the central nose poke, the signal light is either illuminated for signal trials or remains off for non-signal trials. After a 1s delay, the magazines on both sides of the central nose poke are illuminated and the rat can make a response. A reward is delivered if the correct side is chosen. An incorrect decision results in a 5s timeout. If the rat does not respond after a 4s limited hold (LH), the trial ends with an omission scored.

**Fig 1.**
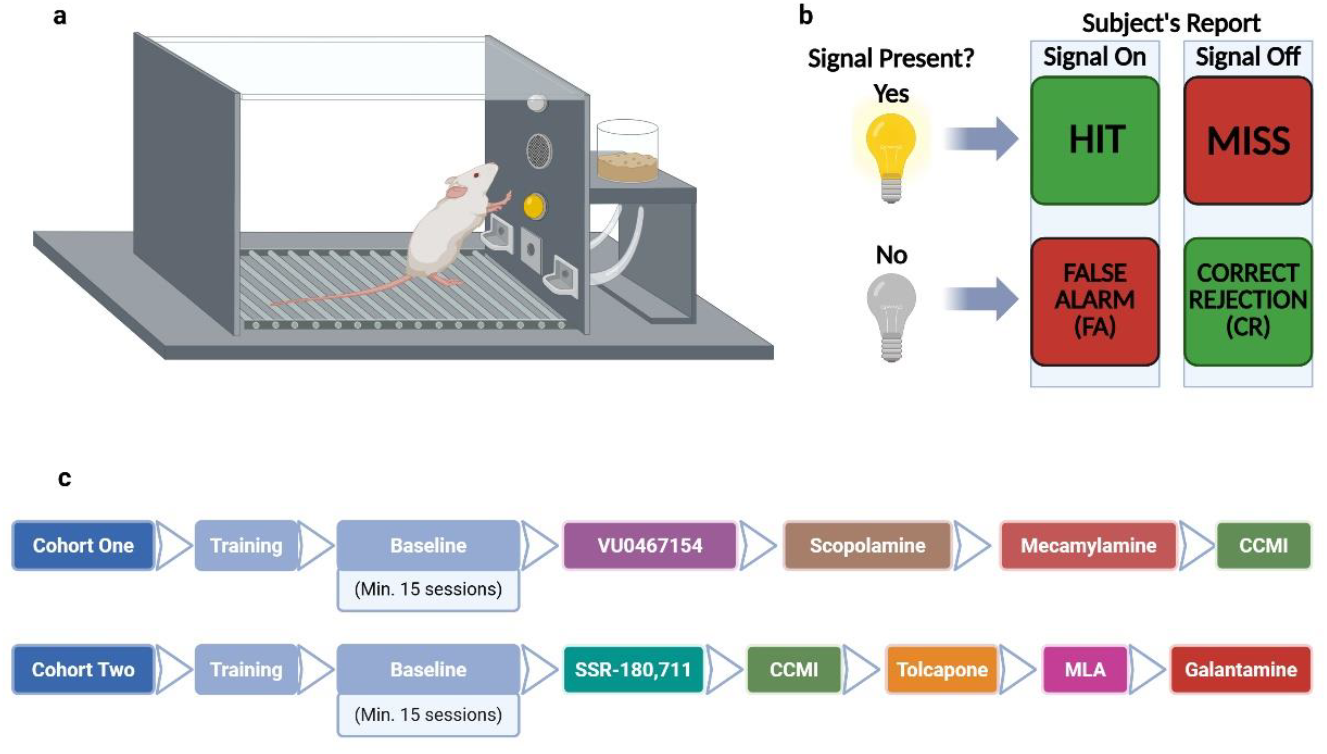
(a) schematic of the operant chamber used for the SDT including a house light, signal light, central nose poke port, and food magazine on each side. The nose poke port and food magazines each contained a light and head entry detector. (b) visual summary of the SDT. (c) experimental timeline. Created in BioRender. Robson, H. (2026) https://BioRender.com/92cdaf4

Premature and otherwise inappropriate head entries and nose pokes were recorded but not punished. Each session consisted of a 20-trial reminder block containing ten 1s signal trials and ten non-signal trials presented pseudo-randomly, followed by a 150-trial test block containing 90 signal trials (signal durations: 0.03s, 0.06s, 0.25s) and 60 non-signal trials presented pseudo-randomly (for further details of SDT protocol, see Wilod Versprille et al. 2026). Following training, all animals were run on the task to a stable level of performance prior to beginning pharmacological interventions. Details of the training stages, experimental timeline, and summary of baseline task performance are shown in the Supplementary Materials. Details of the training stages, experimental timeline, and summary of baseline task performance are shown in the Supplementary Materials.

The task’s design allowed dissociation of specific perceptual and decision-making effects from non-specific changes in motor output. The SDT “go-go” design requires a response on every trial, which ensures that d′ and criterion are not confounded by response omissions. As such, sedative or motor-suppressive drug effects would primarily manifest as changes in response latency rather than shifts in our measures of perceptual and/or decision-making processes.

### Pharmacological interventions

The pharmacological experiments were run on 3-day cycles comprising a baseline SDT session, a drug testing session (SDT with distractors), and a washout day where subjects remained in their home cages. The SDT with distractors ran in the same way as the baseline SDT, except every other 20 trials were distractor trials (making trial 21 the first distractor trial), during which the house light pulsed continuously (1 Hz) from the end of the previous ITI until a response selection had been made. Drug doses were administered in a Latin square design, with a minimum of 48 hours washout between each dose.

### Drugs

To reduce the novelty and potential anxiolytic effects of injections, all animals were given a single habituation injection of sterile saline (1 ml/kg, i.p.) during the final week of training. All final injection solutions were pH-adjusted to 7.0-7.4 and gently warmed where appropriate to reduce viscosity. Dose ranges were determined based on previous literature (e.g., Bushnell et al. 1997; Bubser et al. 2014; Nikiforuk et al. 2015; Nikiforuk et al. 2016; Detrait et al. 2016; Hahn et al. 2011; Kristensen et al. 2007).

### Cohort one

Mecamylamine hydrochloride (Sigma-Aldrich, UK) and scopolamine hydrobromide (Sigma-Aldrich, UK) were dissolved in sterile saline solution. CCMI (MedChemExpress, UK) was dissolved in an aqueous 10% Cremophor solution. An M4 PAM, VU0467154 (Boehringer Ingelheim, Germany) was dissolved in a 20% 2HP-β-cyclodextrin solution. Mecamylamine (0.67, 2, 6 mg/kg), scopolamine (0.03, 0.056, 0.1 mg/kg), and CCMI (0.1, 0.3, 1 mg/kg) and their respective vehicles were administered intraperitoneally at a volume of 1 ml/kg of body weight 20 minutes prior to the start of the task. VU0467154 (0.3, 1, 3 mg/kg) and its vehicle were administered intraperitoneally at a volume of 5 ml/kg of body weight, one hour prior to the start of the task.

### Cohort two

Methyllycaconitine citrate (MLA; MedChemExpress, UK), galantamine hydrobromide (TargetMol Chemicals, UK), and SSR-180,711 (Boehringer Ingelheim, Germany) were dissolved in sterile saline solution. Tolcapone (TargetMol Chemicals, UK) was dissolved in a sterile saline solution with a few drops of Tween-80. CCMI (MedChemExpress, UK) was dissolved in 5% DMSO and 30% PEG300 with a few drops of Tween-80, then suspended in a 40% 2HP-β-cyclodextrin solution. MLA (0.1, 0.3, 1, 3 mg/kg), galantamine (0.3, 1, 3 mg/kg), and CCMI (0.3, 1, 1.8 mg/kg) and their respective vehicles were administered intraperitoneally at a volume of 1 ml/kg of body weight, 15, 30, or 20 minutes prior to the start of the task, respectively. SSR-180,711 (1, 3, 10 mg/kg) and its vehicle were administered intraperitoneally at a volume of 2 ml/kg of body weight, 30 minutes prior to the start of the task. Tolcapone (10, 30 mg/kg) and its vehicle were administered intraperitoneally at a volume of 2.5 ml/kg of body weight, one hour prior to the start of the task.

### Statistical analysis

All analyses were conducted in R (version 4.5.2) and Python (version 3.11.14). Analyses were conducted using all non-omission trials from the 150-trial test block. Within this dataset, each trial yielded a binary report (signal present vs absent) and a binary accuracy outcome (correct vs incorrect).

Sensitivity (d′) and response criterion (c) were estimated within a single probit-link generalised linear mixed model, implemented with the lme4 package with a binomial family, probit-link, and BOBYQA optimiser, referred to here as the SDT-GLMM (Bates et al. 2015; DeCarlo 1998). The subject’s response was coded as −0.5 (signal absent) and +0.5 (signal present), centred on zero with a contrast of 1, such that the intercept estimates -c directly (Macmillan and Creelman 1991). To recover conventional c values (positive c = more conservative bias), intercept-related estimates were negated *post-hoc*. The fixed-effect structure was specified *a priori* to reflect the experimental manipulations and covariates of interest. The random-effects structure included correlated by-animal random intercepts and random slopes for signal presence to model individual variability in overall criterion and d′. The model predicted the probability of each subject reporting signal present or absent as a function of the trial parameters and was specified as: response ∼ signal_c * *dose \** distractor + signal_c:duration_f + sex + signal_c:sex + sex:dose + (signal_c | rat_id) where duration_f is a factor variable of signal duration and signal_c is the encoding of the signal state. The signal_c × dose × distractor interaction tests whether drug-related changes in d′ differ as a function of distractor condition. Correlated by-animal random intercepts and slopes on signal_c allowed the model to account for variability in c and d′ between animals. This approach models each response as the observed outcome of an unobserved decision process.

Although only the report of “signal present” or “signal absent” is measured, SDT assumes that these reports arise from overlapping signal and noise evidence distributions separated by a response criterion. In the probit-link DeCarlo formulation, the signal coefficient estimates the separation between these latent distributions, corresponding to d′, while the intercept estimates response criterion. This allows d′ and bias to be estimated within the trial-level mixed-effects model, rather than calculated *post hoc* from aggregated hit and false alarm rates (DeCarlo 1998).

For comparison to standard metrics of signal detection performance, separate logit-link binomial GLMMs were fitted for hit and correct rejection rates using per-animal aggregated counts: cbind(n_hit, n_miss) ∼ dose * distractor + (1 | subject_key), and equivalently for correct rejections. Hit and correct rejection rates were used rather than general accuracy, as they allow separate assessment of signal and non-signal trials.

To extract parameter values from the SDT and binomial GLMMs, estimated marginal means (criterion) and marginal trends (d′) were computed using the emmeans package (Lenth and Piaskowski 2025). Each dose was compared against vehicle using Dunnett-corrected contrasts, both averaged over distractor and within each distractor level. Contrasts examining whether dose effects differed between distractor conditions were Bonferroni-corrected. The significance level was *p* < .05 for all tests.

## Results

### Task validation

The visual distractor reliably impaired performance across all experiments, confirming task sensitivity to manipulations of attentional load (e.g., for MLA: β = 0.19, SE = 0.04, z = 4.59, *p* < .001). Similarly, shorter signal durations (relative to 0.25 s) reduced performance across all drugs (e.g., MLA: 0.03 s, β = -2.32, SE = 0.06, z = 36.66, *p* < .001; 0.06 s, β = -1.52, SE = 0.06, z = 24.21, *p* < .001). Data are presented collapsed across sex, as no consistent sex effects were observed (see Supplementary Materials for details). Tables 1 and 2 summarise drug effects across outcome measures.

**Table 1.** Summary of drug effects in cohort one. A probit-link GLMM with random intercept and slope per rat was used to evaluate the effects of drug doses on various SDT performance measures. CR = Correct rejection. D+ = Distractor On, D- = Distractor Off. Arrows indicate p-values from Dunnett-corrected comparisons against vehicle: - = p > .1, ↓ or ↑ = p < .1, ↓↓ or ↑↑ = p < .05, ↓↓↓ or ↑↑↑ = p < .01.

| Cohort One | Dose (mg/kg) | Sensitivity ($d'$) | | | Criterion | | | Hit Rate | | | CR Rate | | |
| --- | --- | --- | --- | --- | --- | --- | --- | --- | --- | --- | --- | --- | --- |
|  |  | All | D+ | D- | All | D+ | D- | All | D+ | D- | All | D+ | D- |
| Mecamylamine | 0.67 | - | - | - | ↓ | ↓ | - | - | ↓↓ | - | ↓ | - | - |
|  | 2 | - | - | - | - | ↓ | - | - | ↓ | - | ↓ | - | - |
|  | 6 | ↓↓↓ | ↓↓↓ | ↓↓↓ | ↑↑ | - | ↑ | ↓↓↓ | ↓↓↓ | ↓↓↓ | ↓ | - | - |
| CCMI | 0.1 | - | - | - | - | - | - | - | - | - | - | - | - |
|  | 0.3 | - | - | - | - | - | - | - | - | - | - | - | - |
|  | 1 | - | - | - | - | - | - | - | - | - | - | - | - |
| Scopolamine | 0.03 | - | - | - | ↑↑ | ↑↑↑ | - | ↓↓↓ | ↓↓↓ | - | - | - | - |
|  | 0.056 | - | - | - | ↑↑↑ | ↑↑↑ | ↑↑ | ↓↓↓ | ↓↓↓ | ↓↓ | ↑↑ | ↑↑ | - |
|  | 0.1 | ↓↓ | - | - | ↑↑↑ | ↑↑ | - | ↓↓↓ | ↓↓↓ | ↓ | - | - | - |
| VU0467154 | 0.3 | - | - | - | - | - | - | - | - | - | - | - | - |
|  | 1 | - | - | - | - | - | - | - | - | - | - | - | - |
|  | 3 | - | - | - | - | - | - | - | - | - | - | - | - |

**Table 2.** Summary of drug effects in cohort two. A probit-link GLMM with random intercept and slope per rat was used to evaluate the effects of drug doses on various SDT performance measures. CR = Correct rejection. D+ = Distractor On, D- = Distractor Off. Arrows indicate p-values from Dunnett-corrected comparisons against vehicle: - = p > .1, ↓ or ↑ = p < .1, ↓↓ or ↑↑ = p < .05, ↓↓↓ or ↑↑↑ = p < .01.

| Cohort Two | Dose (mg/kg) | Sensitivity (d') |  |  | Criterion |  |  | Hit Rate |  |  | CR Rate |  |  |
| --- | --- | --- | --- | --- | --- | --- | --- | --- | --- | --- | --- | --- | --- |
|  |  | All | D+ | D- | All | D+ | D- | All | D+ | D- | All | D+ | D- |
| MLA | 0.1 | - | - | - | - | - | - | - | - | - | ↑↑ | - | - |
|  | 0.3 | ↑↑↑ | ↑ | ↑↑ | ↑↑↑ | ↑↑ | ↑↑ | - | - | - | ↑↑↑ | ↑↑↑ | ↑↑↑ |
|  | 1 | ↓ | - | ↑ | - | - | - | - | - | - | ↑↑ | - | ↑↑ |
|  | 3 | - | - | - | - | - | - | - | - | - | ↑↑ | - | - |
| Galantamine | 0.3 | - | - | - | - | - | - | - | - | - | - | - | - |
|  | 1 | - | - | ↑↑ | - | - | - | - | - | - | - | - | ↑↑ |
|  | 3 | ↓↓↓ | ↓↓↓ | ↓↓ | - | - | - | ↓↓↓ | - | ↓↓↓ | ↓ | ↓↓ | - |
| CCMI | 0.3 | - | - | - | - | - | - | - | - | - | - | - | - |
|  | 1 | - | - | - | - | - | - | - | - | - | - | - | - |
|  | 1.8 | - | - | - | ↑↑↑ | ↑↑↑ | ↑↑↑ | ↓↓↓ | ↓↓↓ | ↓↓↓ | ↑↑↑ | ↑ | ↑ |
| SSR-180,711 | 1 | - | - | - | - | - | - | - | - | - | - | - | - |
|  | 3 | - | - | - | - | - | - | - | - | - | - | - | - |
|  | 10 | ↓ | - | ↓↓ | ↑↑↑ | - | ↑↑↑ | ↓↓↓ | - | ↓↓↓ | - | - | - |
| Tolcapone | 10 | - | - | - | - | - | ↑ | - | - | ↓↓ | - | - | - |
|  | 30 | - | - | - | ↓ | ↓↓ | - | - | ↑ | - | - | - | - |

### Non-selective controls

Scopolamine, VU0467154, and tolcapone were investigated to assess whether non-nicotinic cholinergic or catecholaminergic mechanisms contribute to SDT performance. Scopolamine reduced d′ at the highest dose tested (0.1 mg/kg; 95% CI [-0.315, -0.031], *p* = .011), but this occurred alongside profound sedative effects across doses, precluding interpretation as a selective attentional effect. In contrast, neither VU0467154 nor tolcapone produced consistent or interpretable effects across outcome measures. These results are reported in full in the Supplementary Material. The main text results focus on manipulations that produced interpretable changes in signal detection performance.

### Effects of nicotinic receptor antagonism

Nicotinic receptor blockade produced robust impairments in signal detection performance. High-dose mecamylamine produced an effect independent of attentional load by significantly reducing hit rate (95% CI [-0.550, -0.241], *p* < .001; Fig 2b), with a trend to decreasing correct rejection rate (95% CI [-0.348, 0.006], *p* = .062; Fig 2d) for both distractor-on and distractor-off trials. High-dose mecamylamine significantly decreased d′ (95% CI [-0.506, -0.216], *p* < .001; Fig 2a), indicative of a diminished ability to distinguish signal from noise, accounting for the observed decrease in both hit and correct rejection rates.

**Fig 2.**
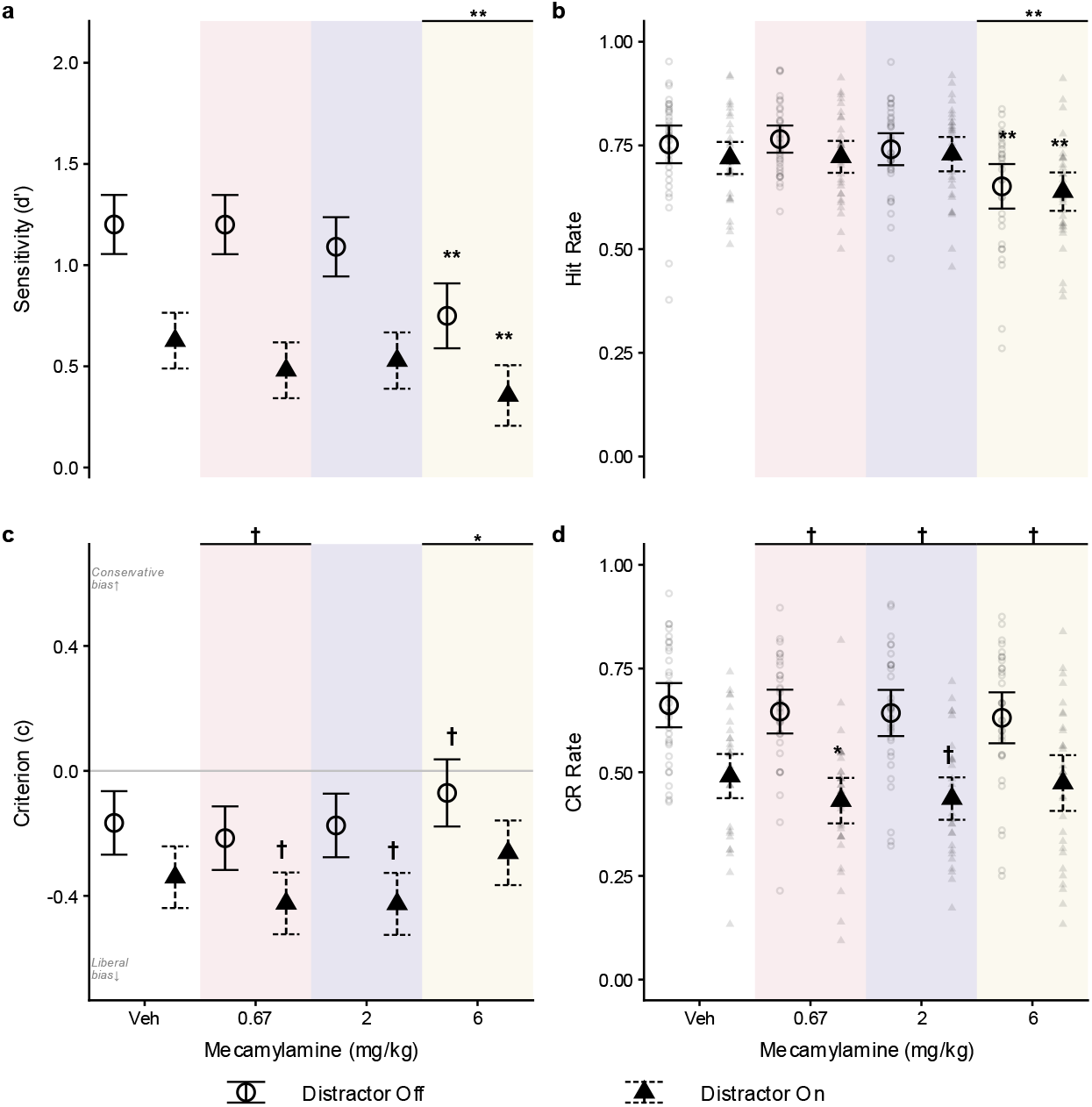
Effects of mecamylamine hydrochloride on SDT performance (n = 31). Left column: signal detection theory parameters estimated from a probit-link GLMM with random intercept and slope per rat. (**a**) Sensitivity (d′), higher values indicate better discrimination. (**b**) Hit Rate. (**c**) Criterion (c), positive or negative values indicate conservative or liberal bias, respectively. Right column: raw accuracy measures. (**d**) Correct rejection (CR) rate. Faded points show individual animal values. For all panels, central markers and error bars show estimated marginal means and 95% confidence intervals. Significance markers indicate Dunnett-corrected comparisons against vehicle. † = p < .1, * = p < .05, ** = p < .01. Veh = Vehicle. Background shading distinguishes dose levels.

### Dissociable effects of acetylcholinesterase inhibition and α7 nAChR antagonism

The acetylcholinesterase inhibitor galantamine produced a dose-dependent improvement in signal detection that was contingent on distractor context. At a moderate dose (1 mg/kg), galantamine improved correct rejection rate (95% CI [0.043, 0.587], *p* = .018; Fig 3d) and d′ (95% CI [0.003, 0.417], *p* = .046; Fig 3a), but these effects were restricted to non-distractor trials, reflected by a significant three-way interaction between galantamine dose x distractor x signal (95% CI [0.034, 0.594], *p* = .022; Fig 3a). At higher doses (3 mg/kg), galantamine impaired performance, reducing hit rate (95% CI [-0.355, -0.058], *p* = .003; Fig 3b), correct rejection rate (95% CI [-0.344, 0.006], *p* = .062; Fig 3d), and d′ (95% CI [-0.381, -0.100], *p* < .001; Fig 3a), but had no effect on criterion (Fig 3c). These findings indicate a context-dependent therapeutic window in which galantamine can facilitate signal detection, but only under low attentional demand.

**Fig 3.**
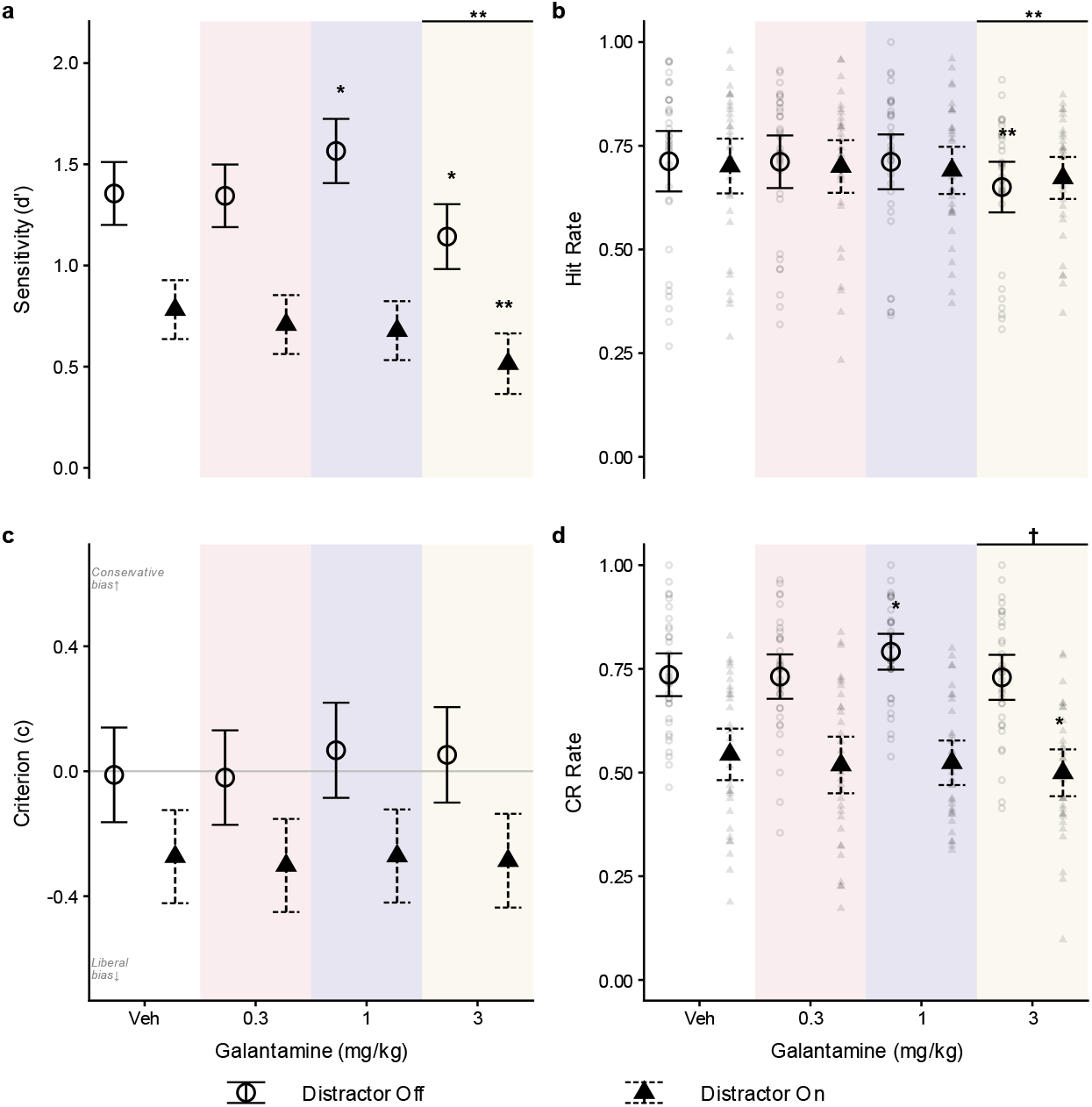
Effects of galantamine hydrobromide on SDT performance (n = 32). For further description of figure elements, refer to Fig 2.

Selective α7 nAChR antagonism with MLA produced robust, dose-dependent improvements in signal detection performance that were dissociable from galantamine improvements. Moderate doses of MLA (0.3 and 1 mg/kg; Fig 4) significantly increased d′ (0.3 mg/kg: 95% CI [0.052, 0.341], *p* = .003; 1 mg/kg: 95% CI [0.001, 0.289], *p* = .047; Fig 4a) and correct rejection rates (0.3 mg/kg: 95% CI [0.169, 0.537], *p* < .001; 1 mg/kg: 95% CI [0.034, 0.399], *p* = .013; Fig 4d), indicating improved discrimination between signal and noise. Uniquely, these effects were relatively robust to distraction (0.3 mg/kg, distractor:ON 95% CI [-0.017, 0.365], *p* = .086; Fig 4a), with a lack of a significant interaction between dose, distractor, and signal status. These distraction-resistant improvements in d′ distinguish MLA from all other compounds tested. The observed improvements following antagonism of α7 nAChRs indicate that reduced α7 signalling enhances signal detection performance in specific contexts.

**Fig 4.**
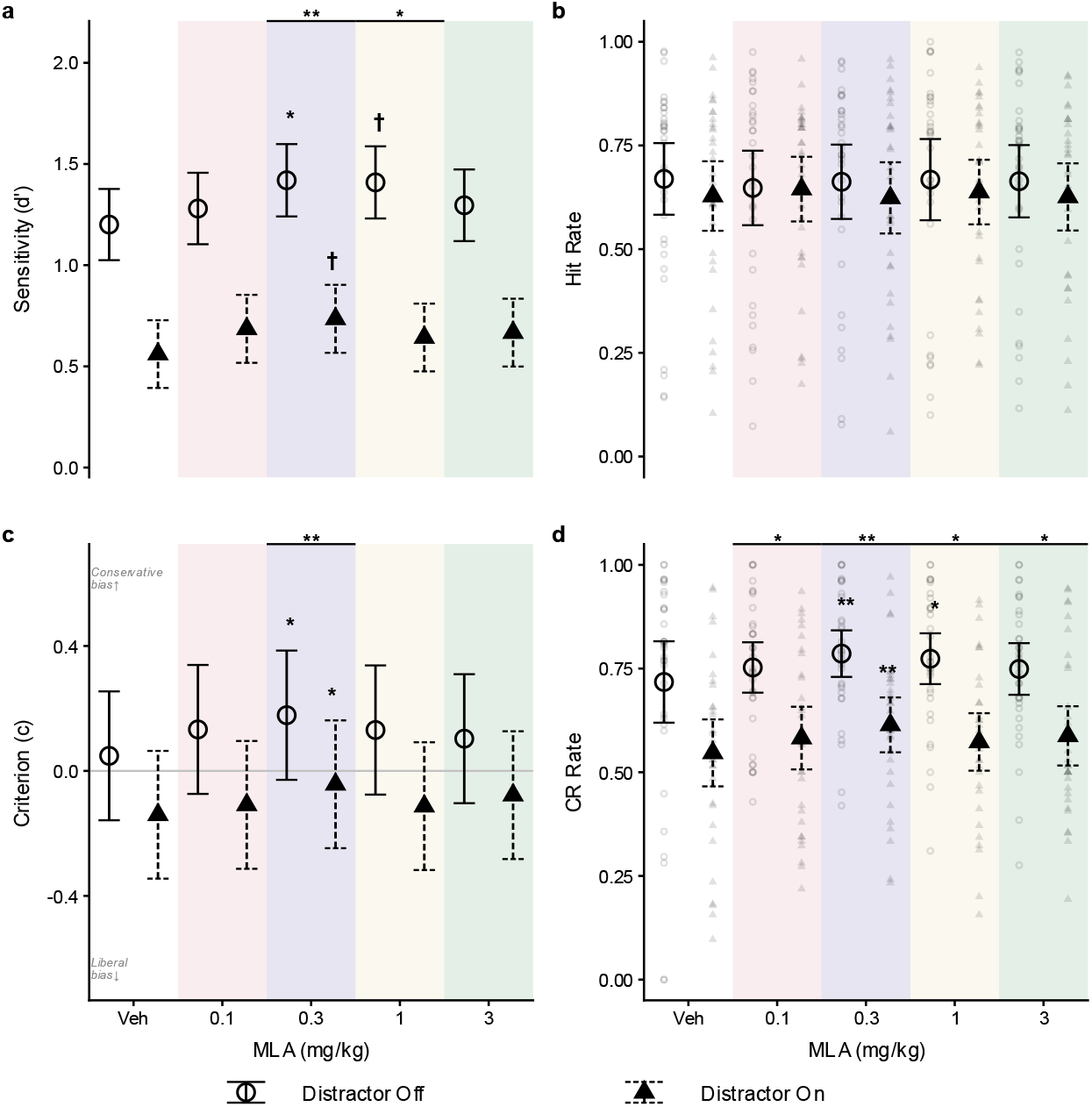
Effects of methyllycaconitine citrate (MLA) on SDT performance (n = 32). For description of figure elements, refer to Fig 2.

At 0.3 mg/kg, MLA also significantly increased criterion (95% CI [0.042, 0.186], *p* < .001; Fig 4c), reflecting a shift toward a more conservative response strategy and indicating that both perceptual and decision processes contributed to performance changes. Conversely, galantamine had no significant effects on criterion across all conditions. These findings suggest that improved performance under moderate MLA reflects both enhanced perceptual sensitivity and, distinctly, a strategic bias toward withholding responses in uncertain conditions.

In summary, MLA produced distraction-resistant improvements in signal detection involving both sensitivity and response strategy, whereas galantamine exerted more limited, context-dependent effects, which primarily influence perceptual sensitivity.

### Impaired signal detection following high dose α7 nAChR activation

High doses of α7 nAChR activation impaired signal detection performance. At the highest doses tested, both α7 nAChR positive allosteric modulation with CCMI (1.8 mg/kg) and α7-partial agonism with SSR-180,711 (10 mg/kg) produced overlapping effects on SDT performance. Specifically, high doses of both compounds impaired hit rates (CCMI: 95% CI [-0.445, -0.159], *p* < .001; SSR-180,711: 95% CI [-0.411, -0.113], p *<* .001; Figs 5b, 6b) and increased the response criterion (CCMI: 95% CI [0.100, 0.234], *p* < .001; SSR-180,711: 95% CI [0.038, 0.176], *p* < .001; Figs 5c, 6c). This shift toward a more conservative response criterion potentially contributed to the observed reduction in hit rates, as subjects became less likely to report the presence of a signal. This increase in conservative bias may also explain the effect of high dose CCMI to increase correct rejection rates (95% CI [0.056, 0.386], *p* = .004; Fig 5d).

**Fig 5.**
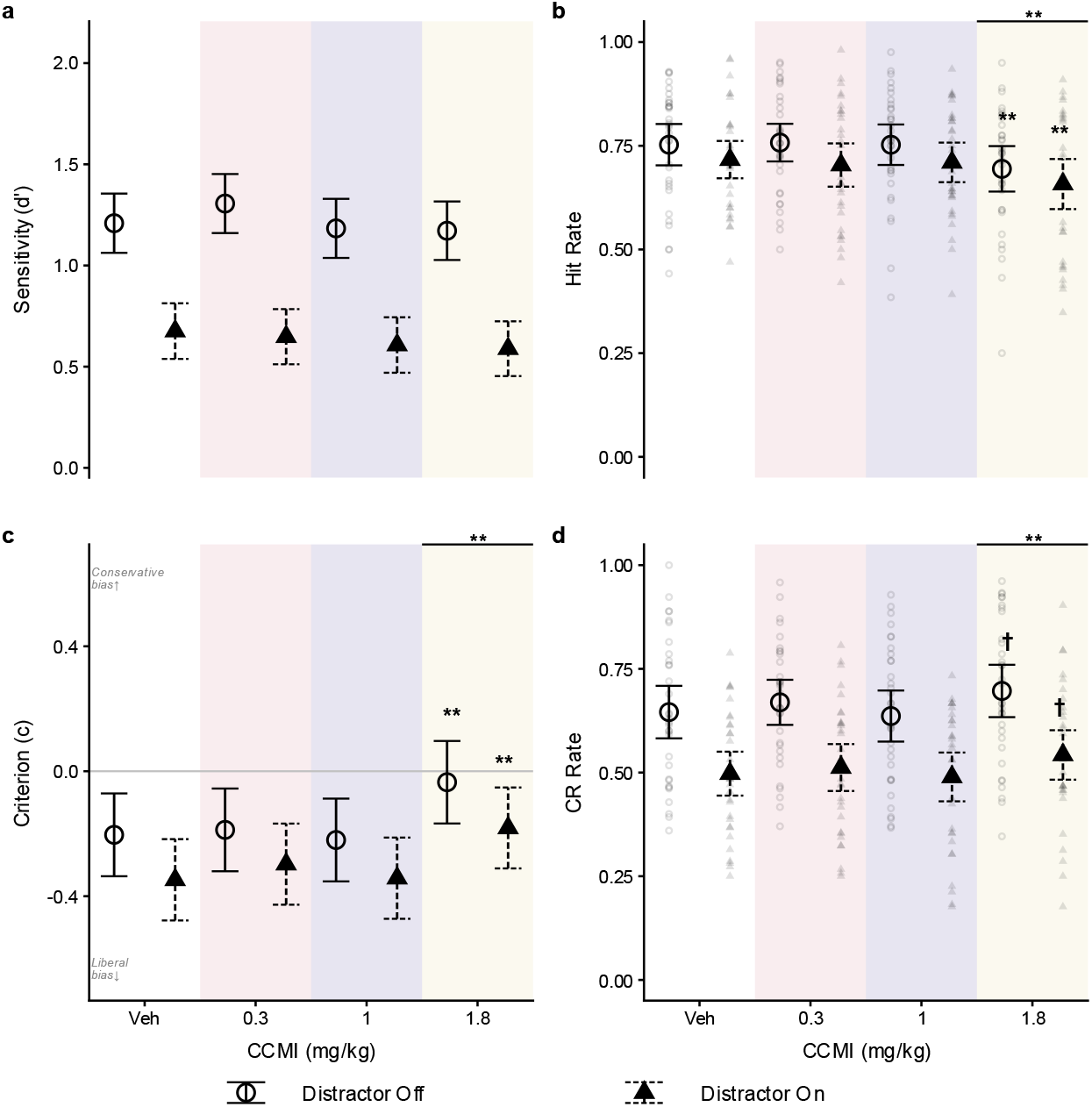
Effects of CCMI on SDT performance measures in cohort two (n = 32). For further description of figure elements, refer to Fig 2.

**Fig 6.**
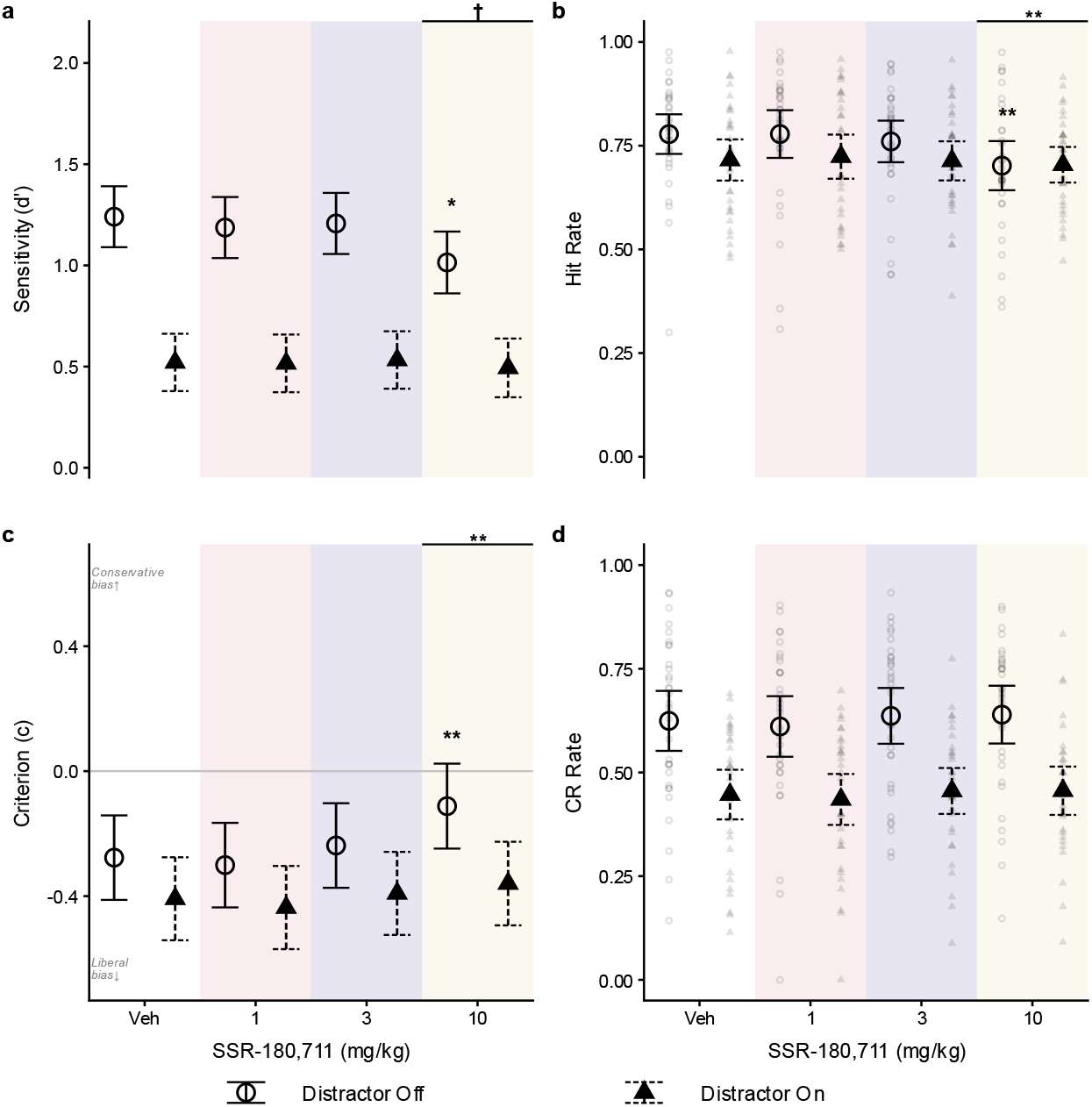
Effects of SSR-180,711 on SDT performance (n = 32). For further description of figure elements, refer to Fig 2.

Unlike CCMI, SSR-180,711 did not significantly affect correct rejection rates (Fig 6d). However, SSR-180,711 significantly decreased d′ during distractor-off trials (95% CI [-0.429, -0.024], *p* = .023; Fig 6a). Thus, the effects of SSR-180,711 share some overlap with those of high-dose galantamine, with impaired hit rates and reduced d′. These convergent findings suggest that galantamine-related impairments may partly reflect excessive α7 nAChR activity.

In summary, pharmacological manipulation of cholinergic signalling produced dissociable effects on signal detection performance. Non-selective nicotinic antagonism impaired d′, whereas acetylcholinesterase inhibition produced moderate, distractor-sensitive improvements. By contrast, selective α7 nAChR antagonism enhanced d′ even under distraction, while α7 nAChR activation impaired performance.

## Discussion

Our findings demonstrate that non-selective nicotinic receptor antagonism, inhibition of acetylcholinesterase, and selective α7 nAChR engagement differentially shape perceptual sensitivity (d′) during signal detection under standard and distractor conditions. Four principal findings emerged. First, non-selective nAChR blockade with mecamylamine reduced d′, suggesting that nicotinic transmission is necessary for maintaining perceptual sensitivity. Second, moderate doses of galantamine, a treatment widely prescribed for cognitive symptoms in AD (Huang and Fu 2010; Marucci et al. 2021), improved d′. However, this benefit was abolished during attentionally demanding conditions (i.e., distraction). In contrast, selective α7 nAChR antagonism with methyllycaconitine (MLA) produced robust, dose-dependent improvements in d′ that were relatively resilient to distraction, distinguishing it from other manipulations. Finally, compounds that positively modulated or directly activated the α7 nAChR impaired signal task performance at high doses. Thus, rather than reflecting a single dimension of ‘cholinergic enhancement’, these findings highlight the importance of receptor-specific and context-dependent cholinergic mechanisms in shaping detection-related processing. Our finding of receptor-specific pro-cognitive effects invites further investigation into the advantages of receptor-specific interventions across different neuromodulatory and cognitive domains.

In the present study, several of the compounds tested produced no clear effects on SDT performance. For example, although scopolamine reduced d′ at the highest dose tested, this effect coincided with marked sedative effects, constraining interpretation of attention-specific effects. Similarly, VU0467154 (a selective M4-AChR PAM) and tolcapone (a COMT inhibitor) had no consistent and interpretable effects on any of the SDT outcomes. Thus, the most robust modulation of SDT performance arose from nicotinic, and particularly α7 nAChR-related mechanisms.

Many pharmacological studies of attention continue to rely on accuracy analysed using ANOVA, limiting mechanistic interpretation. While some studies adopt d′ and criterion (e.g., Wilod Versprille et al. 2026), these are typically derived from aggregated data, reducing power and sensitivity to trial-level variability and individual differences, and limiting flexibility in modelling the influence of experimental covariates. In contrast, we modelled trial-by-trial outcomes within a signal detection theory-based GLMM, capturing individual variability and dissociating drug effects on d′ and response bias across experimental conditions. Using this approach, similar changes in performance were shown to arise from distinct underlying mechanisms, including reductions in d′ (e.g., mecamylamine), shifts in response strategy (e.g., high-dose α7 modulation), or combined effects (e.g., MLA). The trial-by-trial modelling approach was pivotal in identifying and delineating receptor-specific effects.

The present findings support a putative mechanistic framework for signal detection in which successful performance requires phasic ACh transients of sufficient amplitude and appropriate duration to exceed a decision threshold (Gritton et al. 2016; Howe et al. 2010; Parikh et al. 2010; Sarter et al. 2014). Within this framework, mecamylamine may impair cue detection by reducing the effective amplitude of transient ACh signals *via* non-selective nicotinic blockade, preventing decision thresholds from being reached (Parikh et al. 2010). High-dose galantamine increases tonic ACh levels (Hopkins et al. 2012), thereby reducing the phasic-tonic contrast and masking detection-relevant signals, particularly under distraction where tonic ACh load is already elevated (Himmelheber et al. 2000; Passetti et al. 2000; Sarter et al. 2006). In turn, the selective α7 nAChR antagonist, MLA may improve signal detection by altering the temporal dynamics of cholinergic signalling, potentially narrowing the duration of cue-related activity (Parikh et al. 2010). This action may increase the fidelity of cue processing by reducing the influence of temporally diffuse or noise-related signalling. At moderate doses, this would increase the proportion of transients that fall within an optimal duration range while pruning short-duration, noise-related transients (Parikh et al. 2010). However, α7 nAChR antagonism may be maladaptive at higher doses *via* excessive shortening of cue-related cholinergic transients. Excessive positive α7 nAChR engagement (high-dose CCMI, SSR-180,711) may impair successful detection by disrupting the temporal profile of cholinergic signalling, either through desensitisation-related effects or aberrant prolongation of activity (Buccafusco et al. 2009; Papke et al. 2009; Parikh et al. 2010).

Our findings thus suggest that excessive α7 nAChR engagement impairs signal detection performance, whether driven by positive allosteric modulation (CCMI), direct α7 nAChR agonism (SSR-180,711), or elevated synaptic ACh levels (galantamine). Importantly, however, ACh release was not measured directly, and interpretations regarding phasic and tonic signalling are therefore inferential, albeit grounded in a well-established literature (e.g., Parikh et al. 2010; Sarter et al. 2014; Ballinger et al. 2016). Combining signal detection paradigms with real-time measurements of cortical acetylcholine (e.g., Jing et al. 2020) would thus be critical for directly testing the proposed mechanisms.

Mecamylamine produced a robust impairment in d′ at the highest dose tested across standard and distractor conditions, with lower doses showing a graded trend toward impairment. These findings indicate that non-selective nicotinic antagonism disrupts a key process required for signal detection that is not readily compensated for by task familiarity. This is consistent with extensive evidence that intact nicotinic receptor signalling supports thalamocortical transmission, cortical sensory gain, and the amplification of weak or uncertain signals during attentional tasks (Dani and Bertrand 2007; Sarter et al. 2009; Sarter et al. 2014).

Prefrontal nAChRs are essential for evoking and facilitating the transient cholinergic signals that are necessary for cue detection (Parikh et al. 2010). Mechanistically, non-selective nicotinic blockade may reduce the effective impact of cue-evoked cholinergic transients, preventing them from reaching the amplitude threshold required to trigger detection-related responses. Conversely, we would expect nAChR agonists to enhance signal detection by increasing the likelihood that cue-evoked cholinergic transients reach the threshold required to drive detection-related responses.

Two compounds produced improvements in d′, but with distinct context dependence. Under standard conditions, moderate doses of galantamine enhanced d′; however, this benefit was abolished on signal-present trials during distraction. This pattern suggests that AChE inhibition supports attentional performance when tonic cholinergic tone is low, likely by increasing ACh availability and facilitating detection-related signalling. In contrast, the presence of a salient distractor may further elevate extracellular ACh (Himmelheber et al. 2000; Passetti et al. 2000; Sarter et al. 2006), which may obscure temporally precise phasic signalling and reduce the selectivity of signal-related processing. Elevated tonic ACh may also promote receptor desensitisation, further compressing the dynamic range available for phasic responses (Picciotto et al. 2008; Buccafusco et al. 2009; Papke et al. 2009). Our findings suggest that under conditions of high distraction, galantamine fails to enhance, and may even compromise, signal detection performance, highlighting the context-dependent efficacy of cholinergic modulation of attention.

In contrast, selective α7 nAChR antagonism via MLA produced dose-dependent, distractor-resistant improvements in d′ at moderate doses. This dissociation suggests that α7-targeting compounds may overcome limitations associated with current cholinergic treatments (Beinat et al. 2015; Tregellas and Wylie 2019; Magnussen 2025). Mechanistically, α7 nAChRs are well positioned to regulate the balance between signal and background activity (Dani and Bertrand 2007; Koukouli and Maskos 2015), and moderate antagonism may enhance signal-to-noise by reducing background interference. Consistent with this, α7 nAChRs have been implicated in controlling the temporal dynamics of cue-evoked cholinergic transients (Parikh et al. 2010; Wallace and Bertrand 2013). Within this framework, moderate MLA may improve detection by constraining cholinergic signalling within an optimal temporal window, enhancing signal specificity while suppressing noise-like activity. This mechanism may be particularly advantageous under distraction, where increased background activity places greater demands on temporal filtering.

The opposing effects of α7 nAChR antagonism and positive modulation provide converging evidence that α7 nAChRs regulate cholinergic signalling during cue detection. High levels of α7 nAChR engagement impaired performance, potentially *via* receptor desensitisation leading to functional antagonism (Picciotto et al. 2008; Buccafusco et al. 2009; Papke et al. 2009). At the circuit level, excessive α7 nAChR activation may disrupt the temporal profile of cholinergic transients, either by prematurely truncating or excessively prolonging cue-evoked signals (Parikh et al. 2010). In both cases, this would reduce the likelihood that transients fall within the optimal temporal window required for reliable cue detection. Consistent with this notion, high-dose CCMI and SSR-180,711 biased behaviour toward more conservative responding, reflected by changes to correct rejection and hit rates. These findings suggest that effective α7 nAChR-targeting interventions must preserve the temporal precision of cholinergic signalling, as deviations in either direction may compromise signal detection performance.

The present study demonstrates that effective cholinergic enhancement of attention depends on both receptor-specific mechanisms and the environmental context. By dissociating the effects of distinct cholinergic pathways, this work provides a framework for developing more precise and resilient strategies to improve attentional function. Under real-world conditions of competing sensory inputs and fluctuating demands, interventions that preserve the integrity of detection-related processing may be particularly beneficial, as exemplified by the distraction-resistant effects of α7 nAChR antagonism observed here. One potential avenue for future work would be to investigate combination therapies, where current ACh-elevating treatments (e.g., galantamine) are paired with selective α7 nAChR antagonism. A combined approach may enhance detection-related signalling while mitigating the dose-limiting side effects of each compound alone.

## Supporting information

SOM Rates Outputs

SOM SDT Outputs

SOM Training Stages

SOM CCMI C!

SOM Scopolamine

SOM Tolcapone

SOM VU0467154

## Author contributions

HJR, JH and JWD together designed the study; HJR and ARHM conducted the measurements; HJR and ARHM performed data analyses and wrote the first draft of the manuscript, which was edited by JWD, LJFWV and JH; all authors approved the submitted manuscript.

## Declarations

### Funding

This study was partly funded by Boehringer Ingelheim Pharma GmbH & Co. KG, Germany. HJR was funded by a studentship from Boehringer Ingelheim Pharma GmbH & Co. KG, Germany and the Medical Research Council, UK. LJFWV was funded by a Cambridge Trust studentship at Cambridge University, Prins Bernhard Cultuurfonds and VSBfonds from the Netherlands and by an Angharad Dodds John Fellowship at Downing College, Cambridge. ARHM was funded by the Woolf Fisher scholarship at Cambridge.

### Data availability

Data are available upon reasonable request from the corresponding author.

### Ethical approval

Experiments were conducted in accordance with the Animals (Scientific Procedures) Act, 1986, Amendment Regulations (2012) on project licences PA9FBFA9F and PP9536688 following ethical review by the University of Cambridge Animal Welfare and Ethical Review Body (AWERB).

### Competing interests

The authors have no relevant financial or nonfinancial interests to disclose.

### Conflict of interest

No conflicts of interest to declare.

### Open Access

This article is licensed under a Creative Commons Attribution 4.0 International License, which permits use, sharing, adaptation, distribution and reproduction in any medium or format, as long as you give appropriate credit to the original author(s) and the source, provide a link to the Creative Commons licence, and indicate if changes were made. The images or other third-party material in this article are included in the article’s Creative Commons licence, unless indicated otherwise in a credit line to the material. If material is not included in the article’s Creative Commons licence and your intended use is not permitted by statutory regulation or exceeds the permitted use, you will need to obtain permission directly from the copyright holder. To view a copy of this licence, visit http://creativecommons.org/licenses/by/4.0/.

## Notes

### Competing Interest Statement

The authors have declared no competing interest.

