## Supplementary figures and images for "Modulatory effects of α7-nicotinic cholinergic receptors on perceptual sensitivity in a visual signal detection task"

### SOM CCMI C!

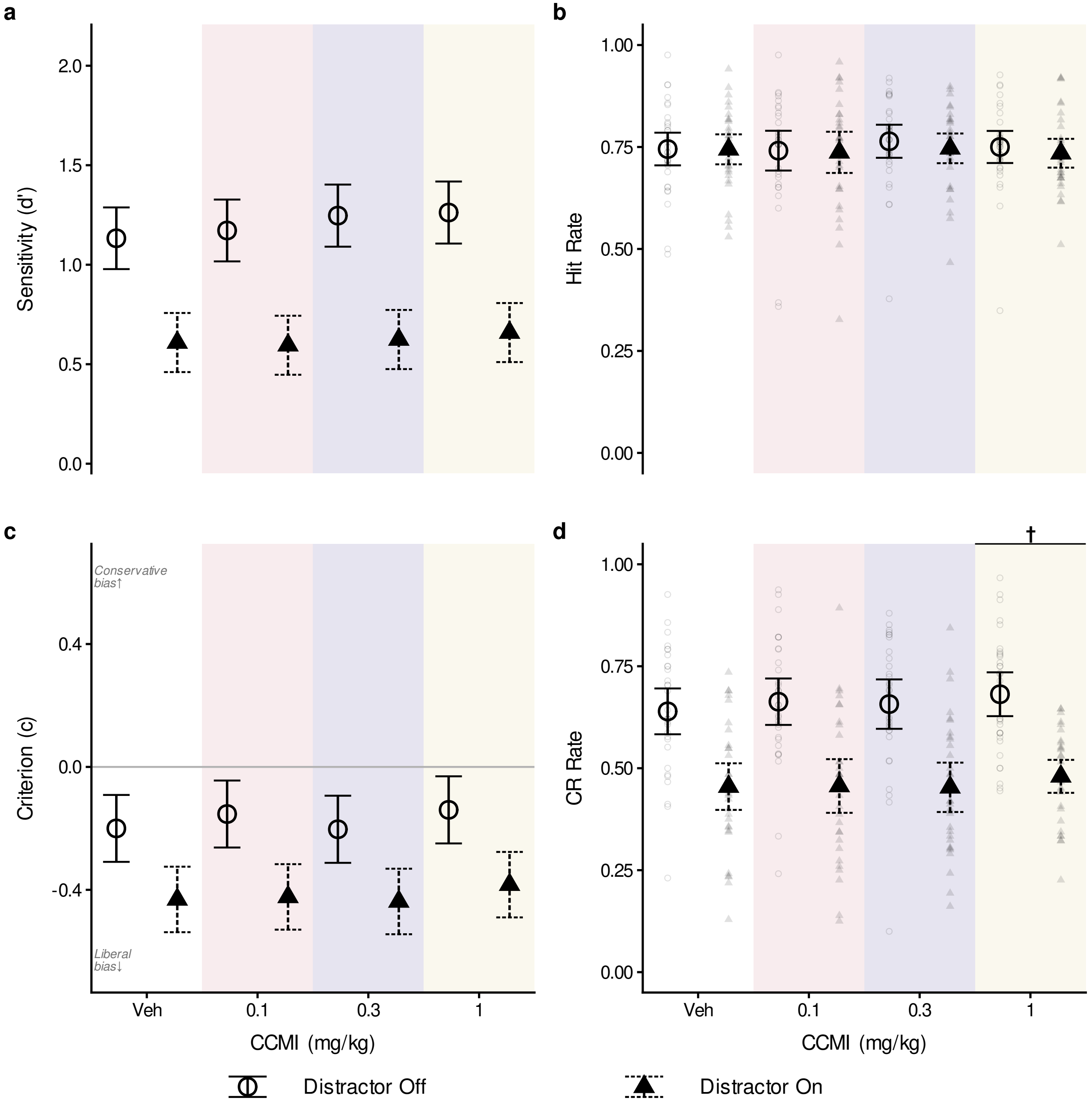

### SOM Scopolamine

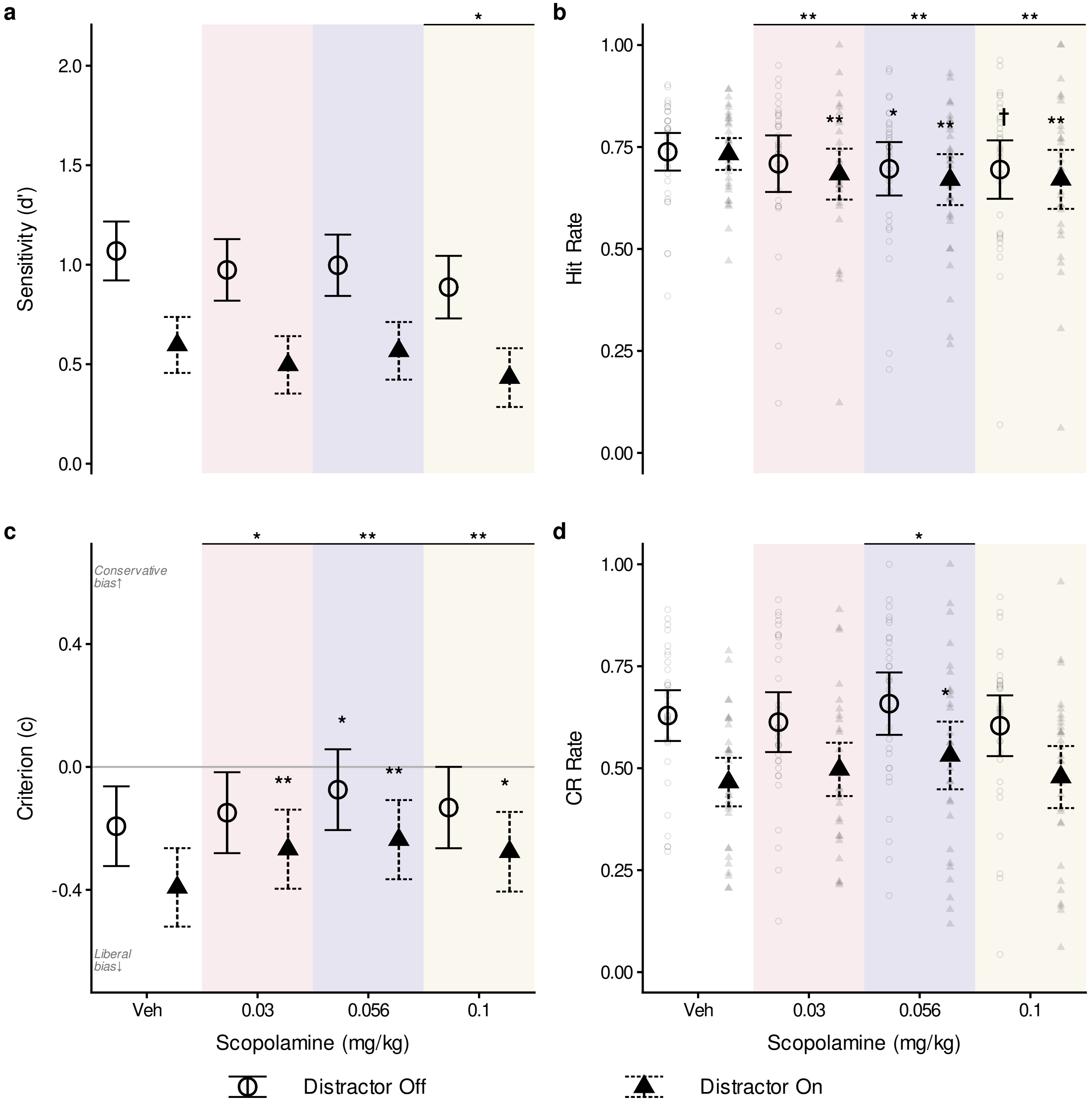

### SOM Tolcapone

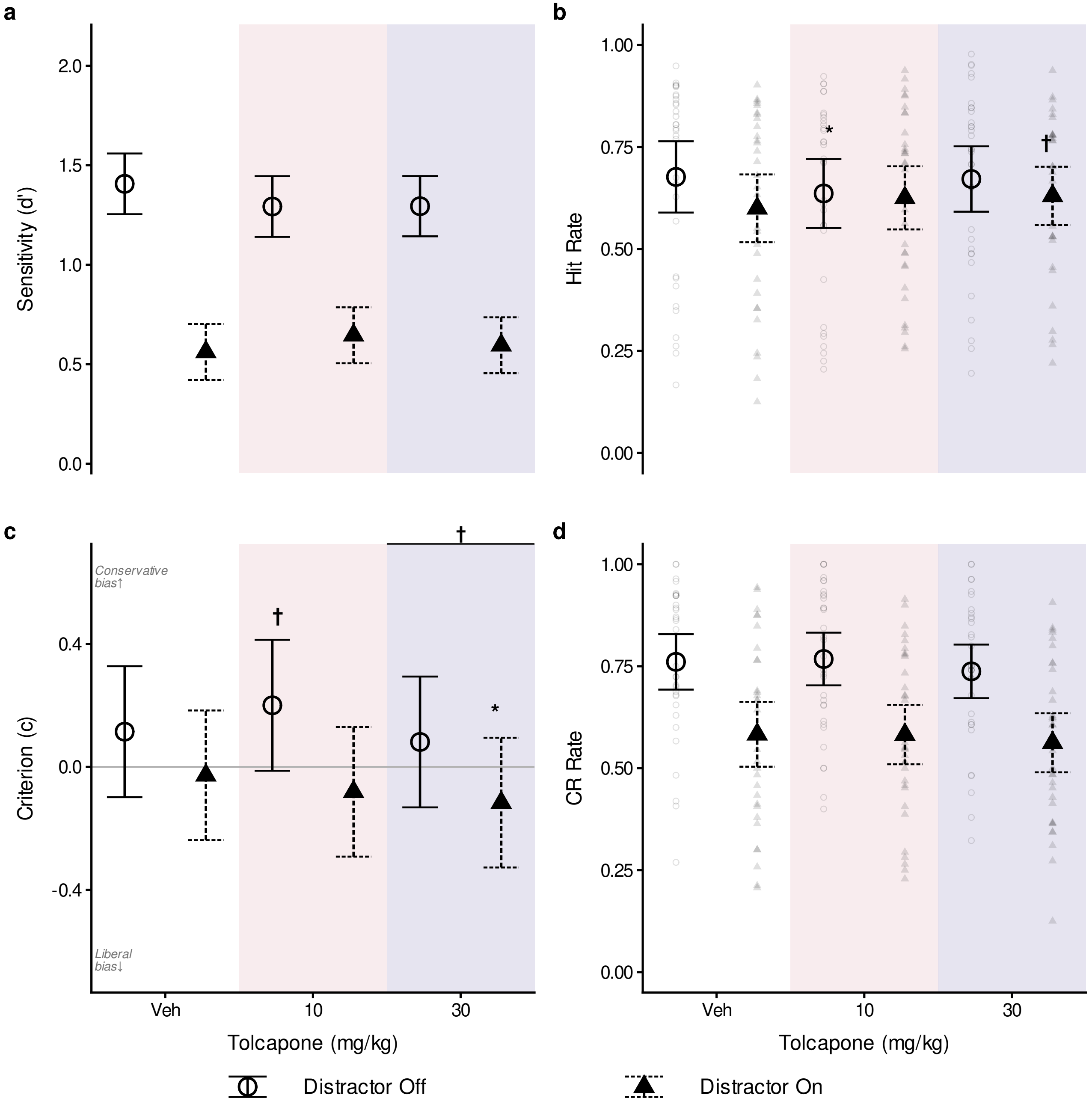

### SOM VU0467154

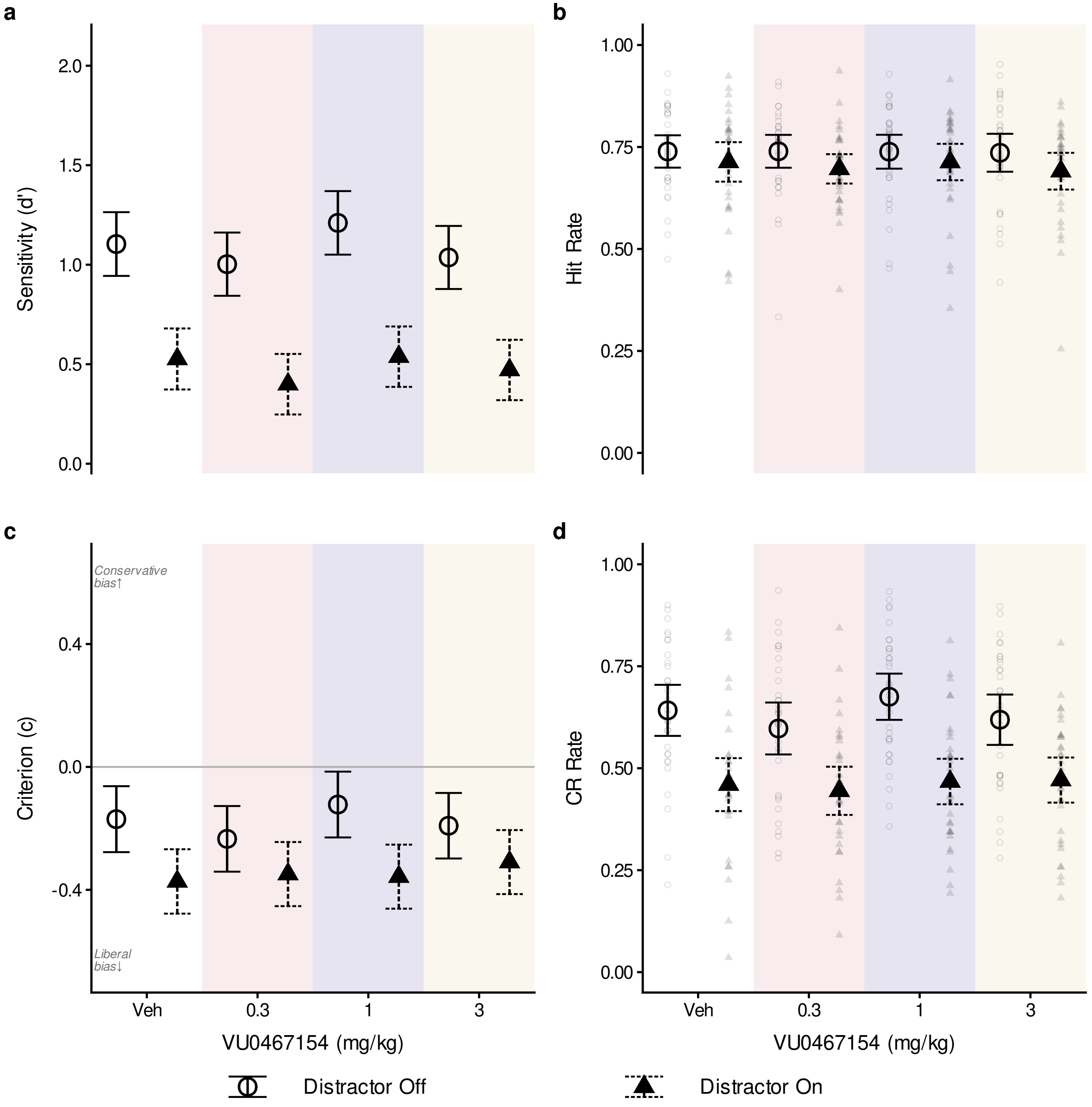
